# Process Optimization of Plasma Bubbling Set Up for Cow Milk Decontamination and Quality Evaluation

**DOI:** 10.1101/2022.12.28.522063

**Authors:** Samarpita Dash, Rangarajan Jaganmohan

**Affiliations:** National Institute of Food Technology Entrepreneurship and Management-Thanjavur, Tamil; Bharathidasan University, Tiruchirappalli, Tamil Nadu, India

**Keywords:** Plasma bubbling, Bovine Milk, GC-MS, Microbiology, Pasteurization, SEM

## Abstract

Cold plasma aims to decontaminate and maintain the naturality of the food. The present study discussed the application of novel plasma bubbling for raw milk pasteurization with different combinations of processing parameters such as voltage, flow rate of air, time interval and volume. The observation showed a decline in microbial load, with an increase in pH value up to 6.9 at optimised conditions (200 volts, the flow rate of air 10 Litres/hour, time 15 minutes and volume 100 mL). The optimized condition was observed was found to be appropriate and could be beneficial to the food industry.

## INTRODUCTION

Milk is an important, widely consumed beverage and crucial to the diet of around six billion of people worldwide as it bestows key macro and micronutrients (Visioli and Strata, 2014). Generally, in dairy-based products, thermal techniques are adopted for enhancing food safety that changes its physicochemical properties including flavour, browning, partial loss of vitamins and depression of freezing point (Topçu et al. 2006). The cold plasma was introduced in dairy processing to ensure food safety and at the same time maintain physicochemical parameters. This non-thermal plasma was successfully used for the decontamination of microorganisms present in meat, milk, water, fresh fruits and vegetables, because of its potential killing effect on microorganisms, such as bacteria, yeasts and fungi. Nowadays it has become a well-known technology in the field of the food industry (Kelly-Wintenberg et al. 1998; Deng et al. 2007; Korachi et al. 2010; Berardinelli et al. 2012).

Cold plasma is generated by both direct and indirect methods. Direct cold plasma application is described by the formation of a large variety of reactive species, with a short life span (~milliseconds). Surface plasma reactions (etching and deposition) was observed in the case of the direct cold plasma method (Misra and Jo, 2017). Indirect cold plasma (remote plasma) generates long-living reactive species such as nitric oxide or ozone that come in contact with the food; while the generation of plasma is done in a separate chamber (Surowsky, et al. 2015). In indirect cold plasma, the quantum of heat transmission to the sample is reduced. So far, an achievement for direct cold plasma has well experimented in the field of food science while the setup for indirect cold plasma is challenging (Misra et al. 2011). This study aims to investigate the effect of indirect dielectric barrier discharge (DBD) i.e., plasma bubbling on the microbial and physicochemical properties of raw cow milk with different process parameters and establishment of an optimisation of plasma bubbling set up for pasteurization of milk.

## MATERIALS AND METHODS

### Plasma bubbling set up

A cold plasma system (plasma bubbling) was developed indigenously. The plasma source was created using a cylindrical aluminium container (plasma generator). In the present study, atmospheric gas was used as feed gas for the generation of free radicals.

Plasma bubbling parameters such as flow rate, voltage and time followed in this study were selected in accordance with the previous studies on decontamination of microorganisms present in milk and coconut neera (Gurol et al. 2012; Aparajhitha and Mahendran, 2019). The cow milk sample was subjected to plasma bubbling treatment with varying voltage (180 and 200 V), flow rate of air at 5 and 10 L/h; the volume of the sample was also varied at 50, 100 and 150 mL, with time intervals of 5, 10 and 15 min (Table 1). The treated samples were compared with the raw cow milk (control sample) for microbial analysis, pH, titratable acidity (TA), colour, total soluble solids (TSS), the morphology of milk powder and gas chromatography-mass spectrometry (GC-MS) analysis.

**Table 1.**
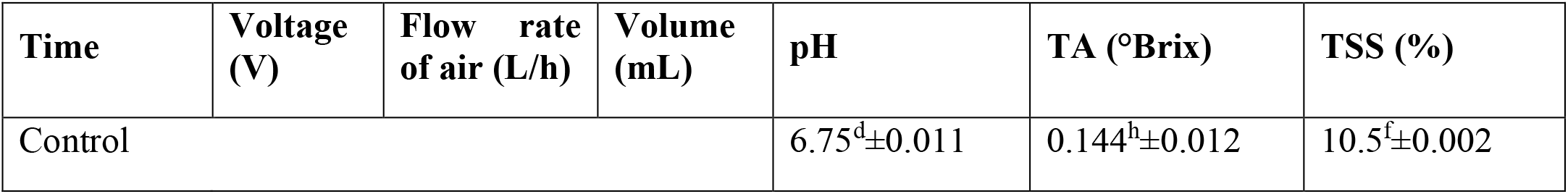
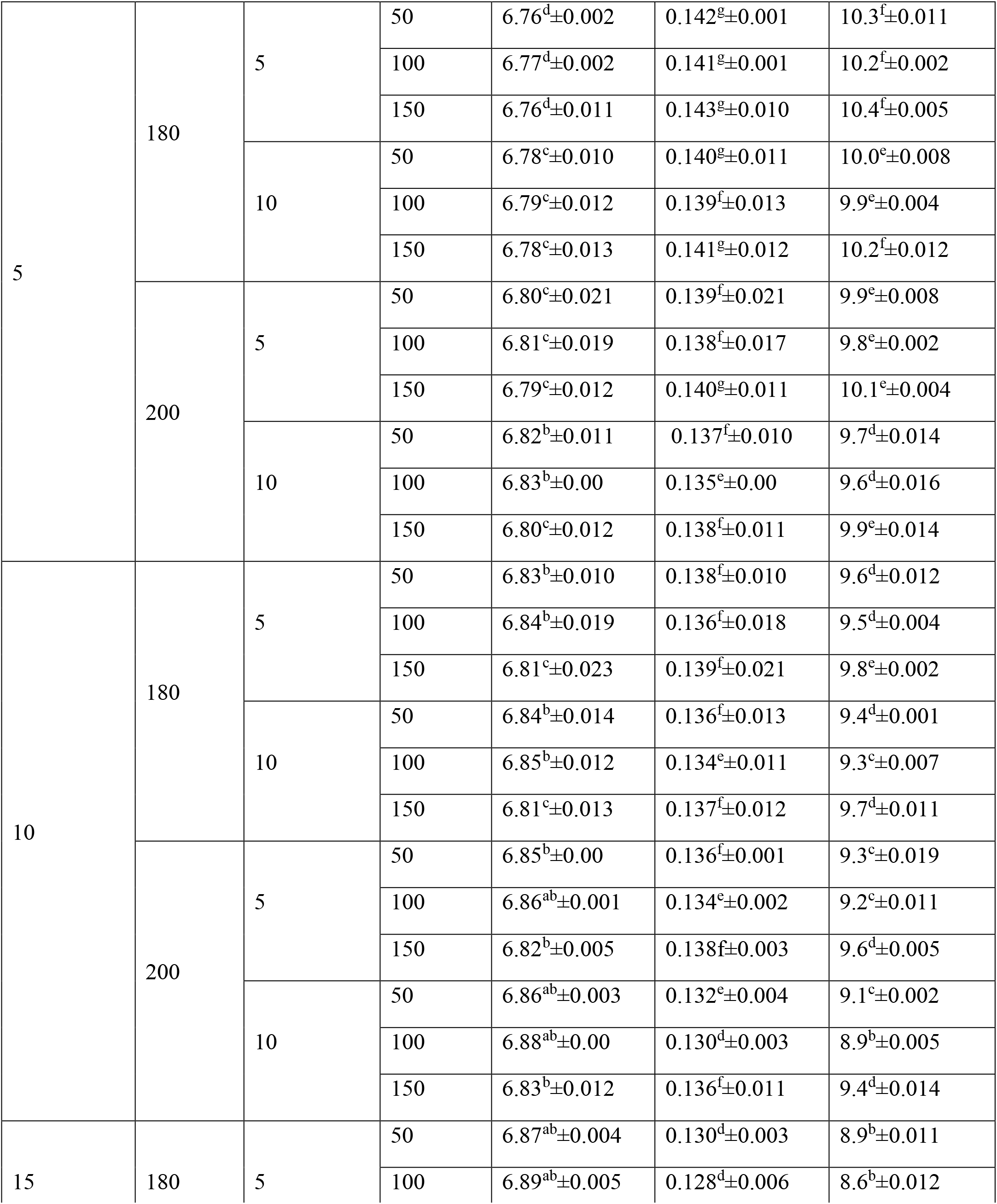
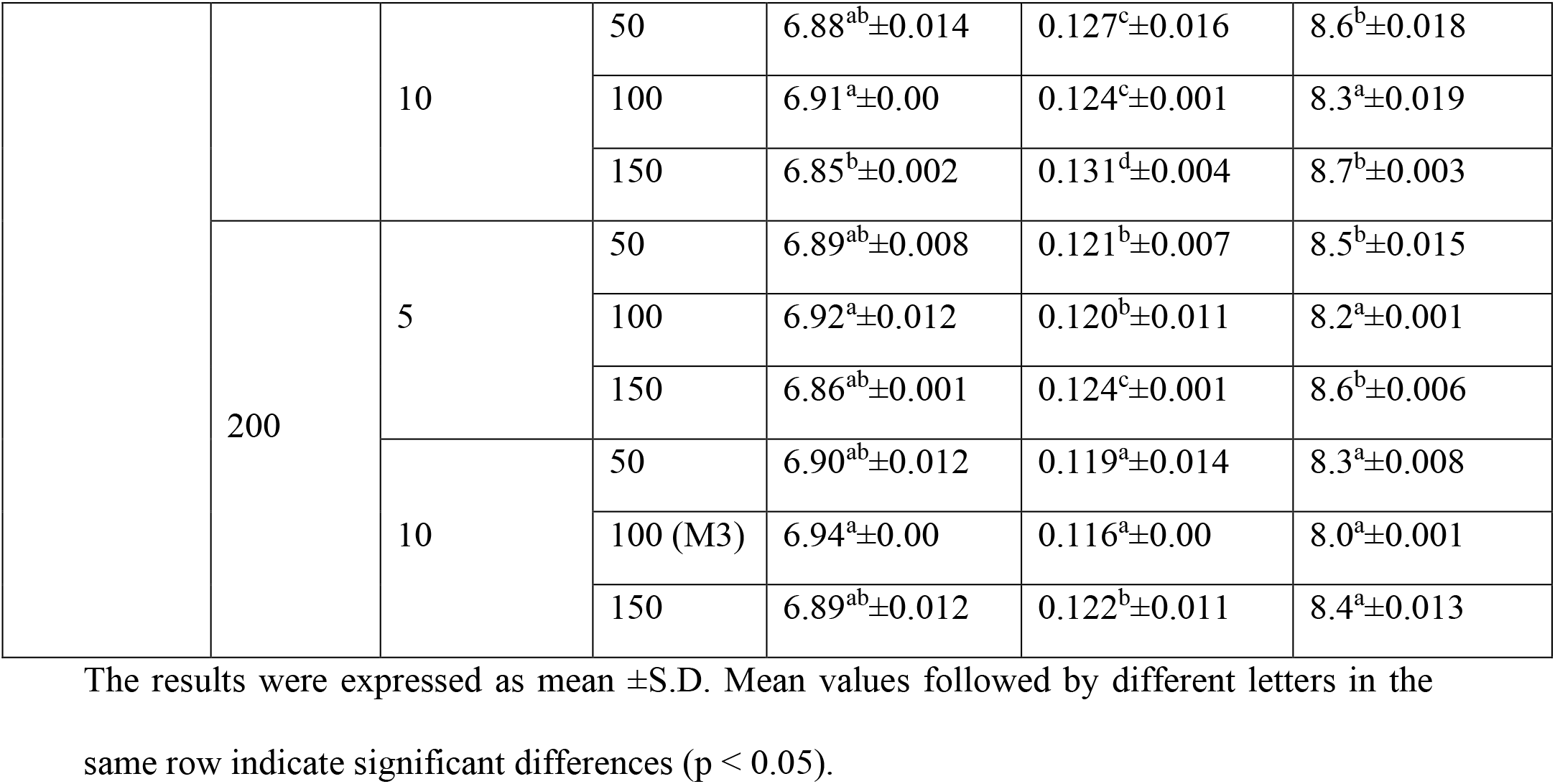
A comparative study on physicochemical properties of control and treated sample for 180 and 200V.

| Time | Voltage (V) | Flow rate of air (L/h) | Volume (mL) | pH | TA ($^\circ$Brix) | TSS (%) |
| --- | --- | --- | --- | --- | --- | --- |
| Control | | | | $6.75^d \pm 0.011$ | $0.144^h \pm 0.012$ | $10.5^f \pm 0.002$ |
| 5 | 180 | 5 | 50 | $6.76^d \pm 0.002$ | $0.142^g \pm 0.001$ | $10.3^f \pm 0.011$ |
| | | | 100 | $6.77^d \pm 0.002$ | $0.141^g \pm 0.001$ | $10.2^f \pm 0.002$ |
| | | | 150 | $6.76^d \pm 0.011$ | $0.143^g \pm 0.010$ | $10.4^f \pm 0.005$ |
| | | 10 | 50 | $6.78^c \pm 0.010$ | $0.140^g \pm 0.011$ | $10.0^e \pm 0.008$ |
| | | | 100 | $6.79^c \pm 0.012$ | $0.139^f \pm 0.013$ | $9.9^e \pm 0.004$ |
| | | | 150 | $6.78^c \pm 0.013$ | $0.141^g \pm 0.012$ | $10.2^f \pm 0.012$ |
| | 200 | 5 | 50 | $6.80^c \pm 0.021$ | $0.139^f \pm 0.021$ | $9.9^e \pm 0.008$ |
| | | | 100 | $6.81^c \pm 0.019$ | $0.138^f \pm 0.017$ | $9.8^e \pm 0.002$ |
| | | | 150 | $6.79^c \pm 0.012$ | $0.140^g \pm 0.011$ | $10.1^e \pm 0.004$ |
| | | 10 | 50 | $6.82^b \pm 0.011$ | $0.137^f \pm 0.010$ | $9.7^d \pm 0.014$ |
| | | | 100 | $6.83^b \pm 0.00$ | $0.135^e \pm 0.00$ | $9.6^d \pm 0.016$ |
| | | | 150 | $6.80^c \pm 0.012$ | $0.138^f \pm 0.011$ | $9.9^e \pm 0.014$ |
| 10 | 180 | 5 | 50 | $6.83^b \pm 0.010$ | $0.138^f \pm 0.010$ | $9.6^d \pm 0.012$ |
| | | | 100 | $6.84^b \pm 0.019$ | $0.136^f \pm 0.018$ | $9.5^d \pm 0.004$ |
| | | | 150 | $6.81^c \pm 0.023$ | $0.139^f \pm 0.021$ | $9.8^e \pm 0.002$ |
| | | 10 | 50 | $6.84^b \pm 0.014$ | $0.136^f \pm 0.013$ | $9.4^d \pm 0.001$ |
| | | | 100 | $6.85^b \pm 0.012$ | $0.134^e \pm 0.011$ | $9.3^c \pm 0.007$ |
| | | | 150 | $6.81^c \pm 0.013$ | $0.137^f \pm 0.012$ | $9.7^d \pm 0.011$ |
| | 200 | 5 | 50 | $6.85^b \pm 0.00$ | $0.136^f \pm 0.001$ | $9.3^c \pm 0.019$ |
| | | | 100 | $6.86^{ab} \pm 0.001$ | $0.134^e \pm 0.002$ | $9.2^c \pm 0.011$ |
| | | | 150 | $6.82^b \pm 0.005$ | $0.138^f \pm 0.003$ | $9.6^d \pm 0.005$ |
| | | 10 | 50 | $6.86^{ab} \pm 0.003$ | $0.132^e \pm 0.004$ | $9.1^c \pm 0.002$ |
| | | | 100 | $6.88^{ab} \pm 0.00$ | $0.130^d \pm 0.003$ | $8.9^b \pm 0.005$ |
| | | | 150 | $6.83^b \pm 0.012$ | $0.136^f \pm 0.011$ | $9.4^d \pm 0.014$ |
| 15 | 180 | 5 | 50 | $6.87^{ab} \pm 0.004$ | $0.130^d \pm 0.003$ | $8.9^b \pm 0.011$ |
| | | | 100 | $6.89^{ab} \pm 0.005$ | $0.128^d \pm 0.006$ | $8.6^b \pm 0.012$ |
| | | | 150 | $6.84^b \pm 0.018$ | $0.134^e \pm 0.018$ | $9.0^c \pm 0.016$ |
|  |  | 10 | 50 | 6.88<sup>ab</sup>±0.014 | 0.127<sup>c</sup>±0.016 | 8.6<sup>b</sup>±0.018 |
|  |  |  | 100 | 6.91<sup>a</sup>±0.00 | 0.124<sup>c</sup>±0.001 | 8.3<sup>a</sup>±0.019 |
|  |  |  | 150 | 6.85<sup>b</sup>±0.002 | 0.131<sup>d</sup>±0.004 | 8.7<sup>b</sup>±0.003 |
|  | 200 | 5 | 50 | 6.89<sup>ab</sup>±0.008 | 0.121<sup>b</sup>±0.007 | 8.5<sup>b</sup>±0.015 |
|  |  |  | 100 | 6.92<sup>a</sup>±0.012 | 0.120<sup>b</sup>±0.011 | 8.2<sup>a</sup>±0.001 |
|  |  |  | 150 | 6.86<sup>ab</sup>±0.001 | 0.124<sup>c</sup>±0.001 | 8.6<sup>b</sup>±0.006 |
|  |  | 10 | 50 | 6.90<sup>ab</sup>±0.012 | 0.119<sup>a</sup>±0.014 | 8.3<sup>a</sup>±0.008 |
|  |  |  | 100 (M3) | 6.94<sup>a</sup>±0.00 | 0.116<sup>a</sup>±0.00 | 8.0<sup>a</sup>±0.001 |
|  |  |  | 150 | 6.89<sup>ab</sup>±0.012 | 0.122<sup>b</sup>±0.011 | 8.4<sup>a</sup>±0.013 |

The results were expressed as mean ±S.D. Mean values followed by different letters in the same row indicate significant differences ( $p < 0.05$ ).

### Microbial analysis

In the current study, the microbes selected for the analysis of raw cow milk, were coliform, yeast and mould. The treated and control milk samples were tested for coliform using violet red bile agar with 10^-4^ and 10^-5^ dilution (Ray and Speck, 1978). For yeast and mould, chloramphenicol yeast glucose agar was used with dilution of 10^-3^ and 10^-4^ (Fleet and Mian, 1987; Torkar and Teger, 2006). Incubation was done using an incubator (CI-10 plus LED Version BOD Incubator, REMI, Madurai, India) at 37°C (for coliform), and 25°C (for yeast, mould). The experimental analysis was performed in triplicate and the result was represented as log CFU/mL (Horáčková et al. 2015).

### pH analysis

The pH of the cow milk sample was analysed by using the pH meter (Laqua PH1100, Horiba Scientific, Singapore) at ambient temperature (Alves et al. 2018).

### Colour measurement

The treated and control samples were poured into cuvette cell (64 mm) and the colour of the sample was evaluated by colourimeter (Colour Flex EZ System, 45/0 LAV); the colour values L* (lightness), a*(redness) and b*(yellowness) were determined (Kathiravan et al. 2014). Before analysis, the instrument was calibrated with standard white and black plates.

### Titratable acidity

The titratable acidity (TA) was performed according to (Muhammad et al. 2019). The result was recorded after observing the endpoint of the light pink colour (Abid et al. 2013) and represented by % of lactic acid as described in (Zakaria et al. 2020).

### Total soluble solids

The total soluble solids (TSS) content was measured at room temperature (22±2 °C) by a digital refractometer (Erma, Japan, 0-80° Brix). The refractometer was calibrated with distilled water before measurement of TSS value and the result was represented as Brix° (Cavalcanti et al. 2008).

### Samples Preparation

The GC-MS analysis was conducted with the milk obtained from optimised conditions (200 V, 10 L/h, 15 min, 100 mL), due to the appropriate decontamination of microbial load after exposure to plasma bubbling, and was compared with the control sample. 10 mL of cow milk (both control and treated) was subjected to fat extraction according to the method reported by (Stefanov, et al. 2010). The analysis of FAME was done (Firestone, 2009).

### GC-MS analysis

A sample volume of 2μL has injected in split less mode until 1 min then split 10:1. Helium was used as carrier gas at a flow rate of 1mL/min. The separation was performed on a column Rtx-5MS (5% Diphenyl / 95% Dimethylpolysiloxane), 30m x 0.25mm ID x 0.25μm df supplied by GC-MS Trace 1310-ac /MS TSQ-900 (Thermo Scientific, USA). The oven program was as follows: initial temperature 110°C for 3.50 min, increase up to 200°C at a rate of 10°C/min, increase to 280°C at a rate of 5°C/min, which was held for 12 min, equilibration time 1 min. For the detection of MS, the mass range was fixed from 50-500 amu, MS source temperature was 230°C and MS Quad temperature was 250°C. Electron ionization was performed with 70 eV. The total run time was 40.50 min for interval samples.

### Statistical analysis

The result was analysed using SPSS version 22 (SPSS, Inc., United States). All the experimental analyses were replicated three times. The statistical analysis was performed using a one-way analysis of variance (ANOVA). The significant differences among the mean values were determined by Duncan’s multiple comparison tests at a confidence level of p<0.05.

## RESULTS

### Microbial analysis

#### Coliform count

A notably decrease in the coliform count was observed after exposure to plasma bubbling. While a decline of cell count was noticed at a voltage of 200 V, the flow rate of air (10 L/h), 100 mL of the volume of cow milk at 15 min treatment time in the plasma bubbling system. Generally, the acceptable range of coliform in milk is 0-1000/ mL at 24 h with 37°C (ISO, 2006) which was obtained by the plasma bubbling treated sample at 200 V, 10 L/h, 15 min and 100 mL of sample volume. Bubbling of plasma treatment on cow milk at voltage 200 V, 10 L/h flow rate, 15 min of treatment time and 100 mL of the sample was able to reduce the population of food-borne pathogens such as coliform, yeast and mould to not detectable levels; hence considered as the optimised condition of the plasma bubbling set up. The reduction observed could be explained due to the production of hydroxyl radicals and oxygen during cold plasma, which damages the cell wall (Surowsky et al. 2014). Again, when microorganisms are exposed to high bombardment to plasma radicals such as •OH and •NO, the bacterial surface absorbed the radicals produced by plasma and create volatile compounds like CO_2_ and H_2_O. Thus, cell death happened as these compounds damage the cellular surface, the repair of the cell surface is not possible (Misra and Jo, 2017). Phan et al., 2017 reported that the interaction of RNS and ROS, on the double bond of the lipid bilayer in the microbial cell, made a strong oxidative effect which was responsible for the damage of transportation of macromolecules inside and outside of the cell (Phan et al. 2017); thus, can lead to pathogen inactivation. An extensive comparison of the coliform decontamination of cow milk has been presented in (Fig 1).

**Fig1.**
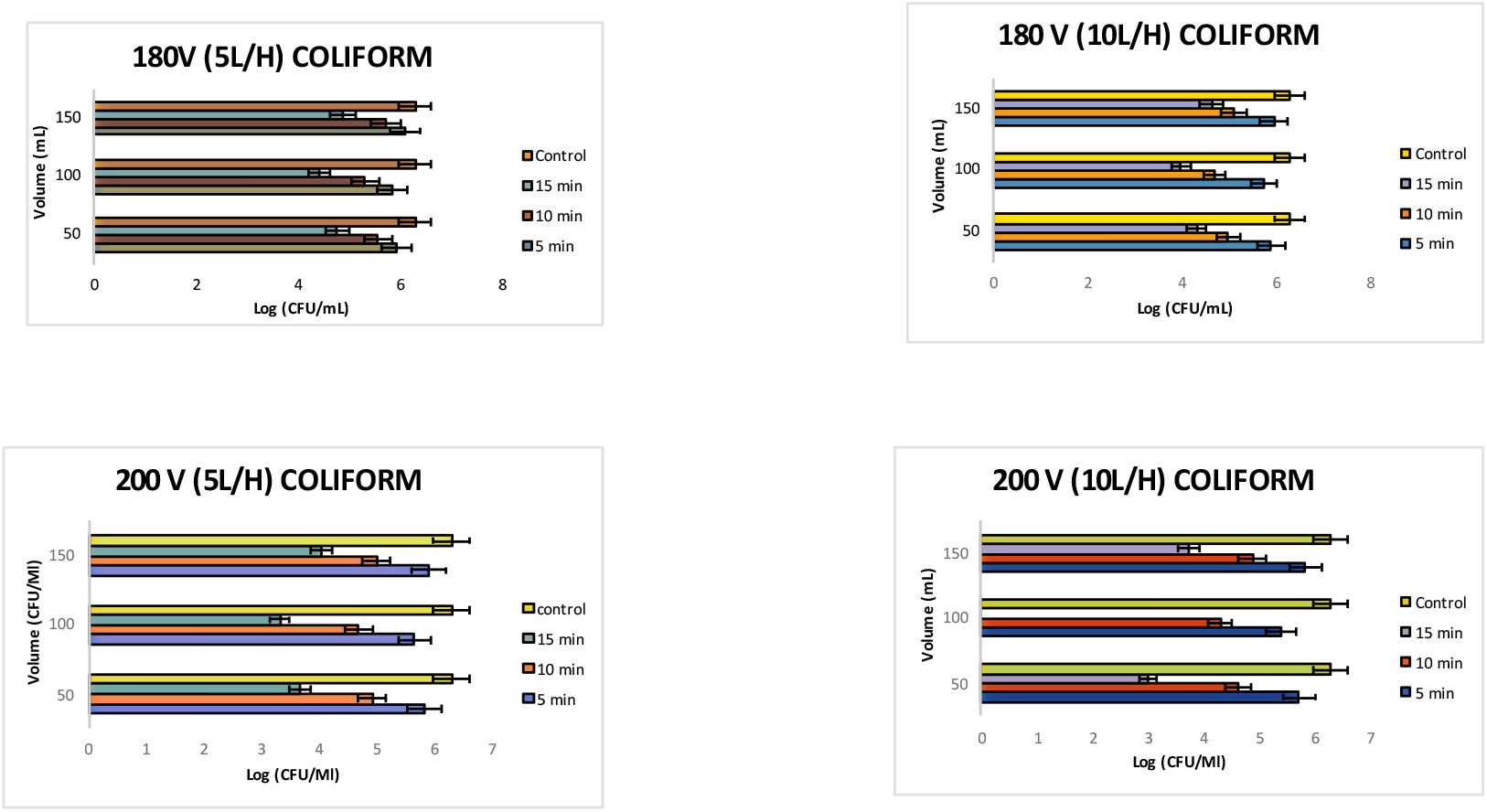
A Comparative study of Viable cell count for coliform for treated and control sample.

**Fig 2.**
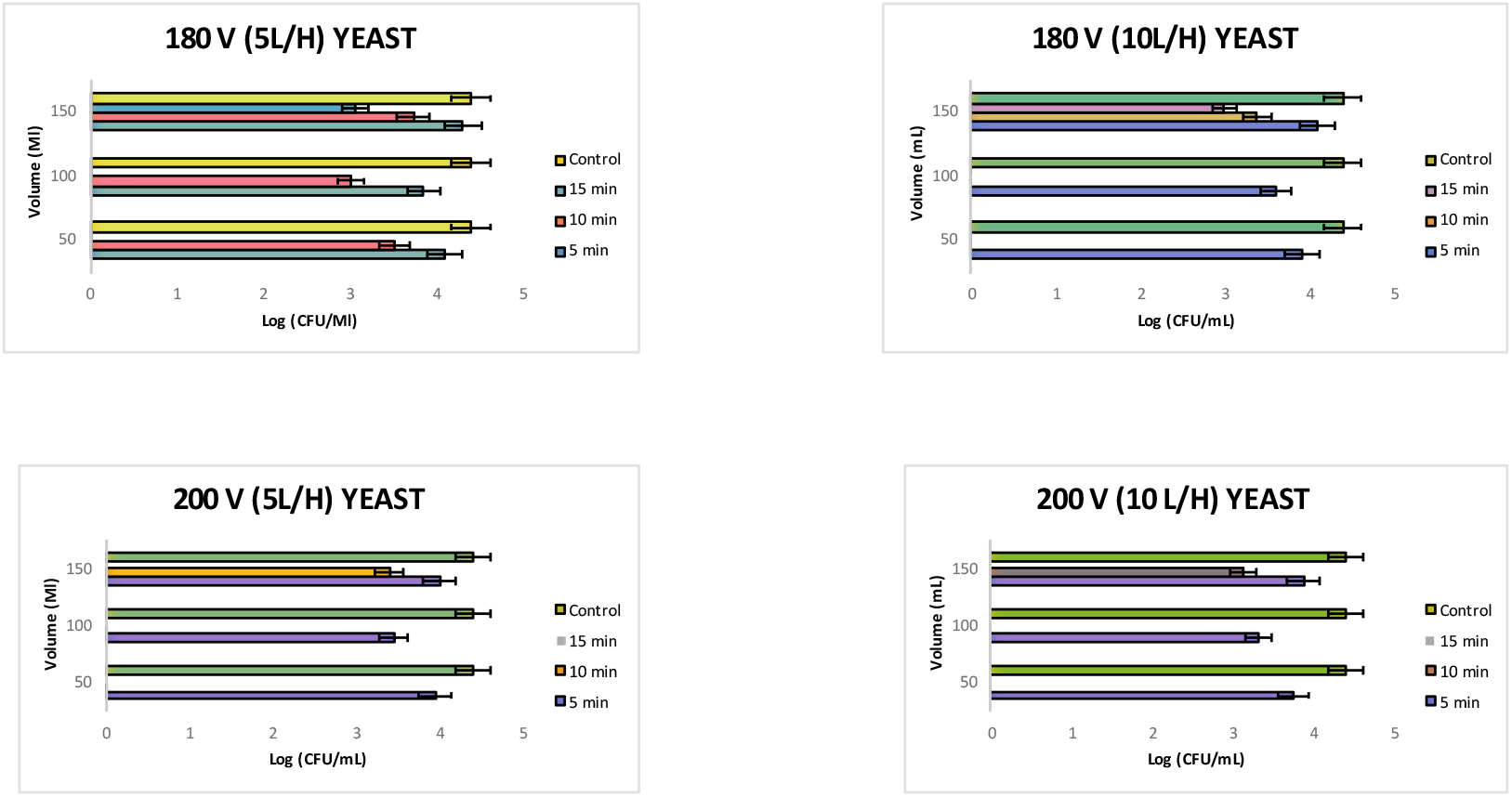
Viable cell count of yeast for treated and control sample.

**Fig 3.**
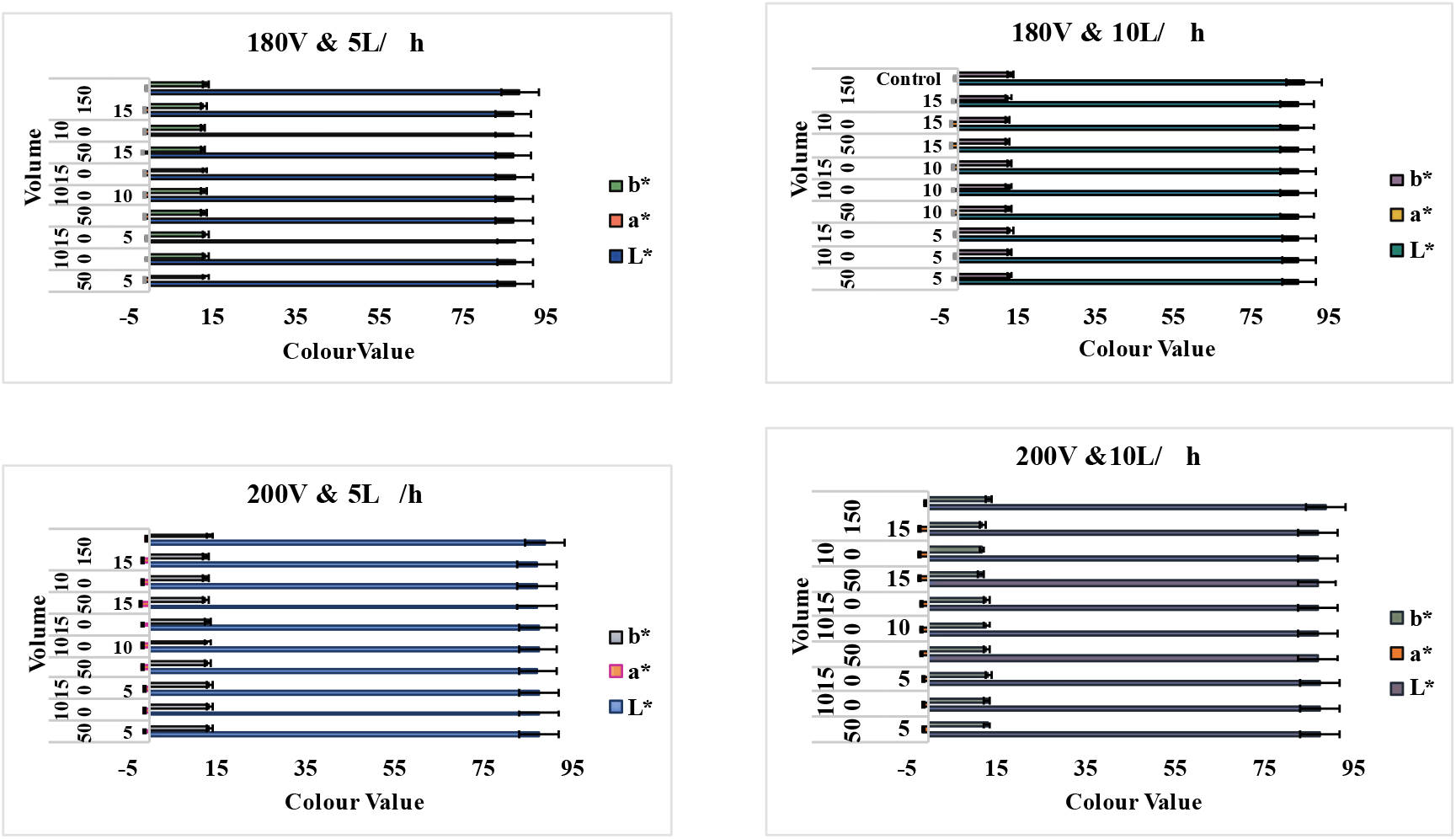
Colour value of Treated and Control sample.

In the present study, the 100 mL of sample volume showed better microbial reduction compared to 50 mL and 150mL could be due to the surface area of the liquid. According to the previous report by Song et al., 2009, the volume of food samples plays a major role in the microbial deactivation on the surface of the food material as reactive plasma species can spike into food to a minimal extent (Song et al. 2009). Many studies have concentrated on the decontamination of dairy foods. However, the mechanism of cell death is still poorly understood (Gurol et al. 2012; Coutinho et al. 2018; Wu et al. 2021).

#### Yeast and mould

7 The control sample of yeast count was 4.4 log CFU/mL while for mould the control value was 5 log CFU/mL. Among the tested organisms, mould proved to be the most sensitive to plasma bubbling treatment, since there was no growth of mould observed after plasma bubbling of the milk. Whereas it was observed that at particular time intervals as well as voltage and flow rate have a major impact on the yeast cell death, yeast cells required particular dosages of plasma bubbling parameters to obtain an optimum reduction. This observation for yeast cells could be due to a different kind of stress produced by the cold plasma generation such as reactive oxygen and nitrogen species which are responsible for the damage of cellular components of yeast cells, as a result, cell death was observed (Polčic and Machala, 2021). Generally, the yeast cells are found in both raw and pasteurised milk (Foster et al. 1957; Randolph et al. 1973; JONES and LANGLOIS, 1977; Fleet and Mian, 1987) but at a low population mostly 10^3^ cells /mL (Fröhlich-Wyder, 2003), while in the present study the input parameters of plasma bubbling treated sample markedly affect the decontamination of yeast cells which indicates a good finding.

Further, a drastic reduction was observed for mould cells after exposure to the plasma bubbling system could be explained due to the breakage of vesicle and the conidiophores after exposure to cold plasma which leads to cell leakage and loss of viability; in this way, cold plasma damages the fungal cell (Suhem et al. 2013). This also may be because indirect plasma contains longer living reactive species like nitric oxide which comes in contact with food materials and kill mould (Surowsky, et al. 2015; Strohm et al. 2019). The overall finding of our result correlates with Coutinho et al. 2018 according to the study, dairy product when exposed to non-thermal plasma treatment, the microbial load in it depends on different factors such as target species of microorganism, the interval of time used for treatment, input power and gas and food composition (Coutinho et al. 2018).

#### pH analysis

A significant increase in pH value was observed after exposure to plasma bubbling. The maximum value of pH was observed at 200 V,10 L/h, 100 mL, 15 min i.e., of 6.94±0.00, while the pH value for the control sample was 6.75±0.011 (Table 1). A previous report on plasma bubbling showed that an increase in OH• radical by increasing time and flow rate observed by electron paramagnetic resonance (EPR) (Aparajhitha and Mahendran, 2019). Another study revealed that indirect DBD plasma contains longer-living reactive species such as nitric oxide or ozone (Surowsky, et al. 2015).

#### Titratable acidity

After exposure to plasma bubbling, a significant decrease in % lactic acid value was observed as compared to the control sample. The (200V,10L/h,100mL,15min) had a value of 0.116±0.00% w/v lactic acid whereas the control sample value of lactic acid was 0.144±0.012% w/v (Table 1). This could be explained due to the increase of OH^•^ radicals formed by the decomposition of an additional water molecule during cold plasma (Guo et al. 2015). Another report explained by (Aparajhitha and Mahendran, 2019) that plasma bubbling produced OH• radical. In this study, with an increase of time, voltage and flow rate of air the OH• radical also increases.

#### Total soluble solids

A decreased value of TSS was observed after a long time of exposure to plasma bubbling at 200V, 10 L/h, 100mL, 15 min i.e., 8° Brix; while the value for control was ~10.5° Brix (Table 1). The decrease can be postulated as due to the ozonolysis, which occurred during plasma exposure thus forming cleavage of the glycosidic bond and helping macromolecule to de-polymerize (Ben’Ko, et al. 2013; Almeida et al. 2017).

#### Colour

A significant decrease was observed for L*, a* and b* values (Fig 4) could be explained due to the degradation of pigment partially after exposure to cold plasma (Lacombe et al. 2015; Ramazzina et al. 2015). As per the experiment, the redness of colour a* value was highest in the case of control cow milk i.e., (−0.96±0.012), whereas, the lowest a* value had observed in the case of (200V,10L/h, 50mL and 15 min) of plasma bubbled milk (−1.21±0.001) indicating slight increase towards green colour. Regarding yellowness (b*coordinate), the lowest value had 13.09 ±0.029 plasma bubbled (200V,10L/h, 50mL and 15 min) while the highest value of b* had the control sample 13.56 ±0.013. Further, the control value of cow milk was brighter than the plasma bubbled cow milk i.e, L* value. Moreover, the carotenoid is indicating the yellow colour of milk and is responsible for the sensorial characteristics of the product (Barrefors *et al*., 1995). Whereas, it was observed that carotenoid content was decreased after exposure to cold plasma treatment with an increase in time (Fernandes, Santos and Rodrigues, 2019). In this study, at 50 mL of sample volume more decrease in colour was observed. The observation found that the volume of the sample plays a crucial role in colour change. The lowest volume of sample was found to be more different in colour. However, the changes in the colour value of milk were not observed in the naked eye through a decrease in colour was noticed. A similar observation on colour value detection by the naked eye was reported on fresh cheese (Wan et al. 2021). According to the previous study, longer treatment exposure of 20 min leads slight change in colour with a ΔE value of 0.52 (Gurol et al. 2012); further, (Kim et al. 2015) suggested that a slight colour difference was noticed in post plasma treatment.

#### GC-MS

The fatty acid composition of the cow milk is presented in (Tables 2 and 3). A total of 12 fatty acids were detected in the control as well as plasma-treated sample, including the saturated chain fatty acids (SFAs, C8:0 - C18:0) and unsaturated fatty acids (MUFAs, C16:1c9, C16:1c7 and C18:1c11). The major fatty acid identified in the samples were palmitic acid, palmitoleic acid and methyl stearate in the case of the control sample with a peak area of 40.63 %, 21.54% and 16.08% respectively. While in the case of plasma bubbling treated at (200V, 10L/h, 100mL, 15 min: PT1) sample; the major fatty acid was palmitic acid, palmitoleic acid, tridecylic acid, methyl stearate with a peak area of 40.53%, 14.25%,12.84% and 11.54 % respectively (Table 3), which could be due to at high voltage of cold plasma, thus leading to degradation and modification of compound (Shi et al. 2017). Again, another report by sarangapani et al. 2017, that the formation of hydroxyl radical during cold plasma treatment may cleave the double bond of unsaturated fatty acid, leading to oxidation (Sarangapani et al. 2017). A similar observation of degradation and modification of fatty was found in cold plasma treated milk (Korachi et al. 2015). Whereas the formation of undecanoic acid found in this study which is having antifungal characteristics along with synergistic effect (Mai et al. 2021).

**Table 2.**
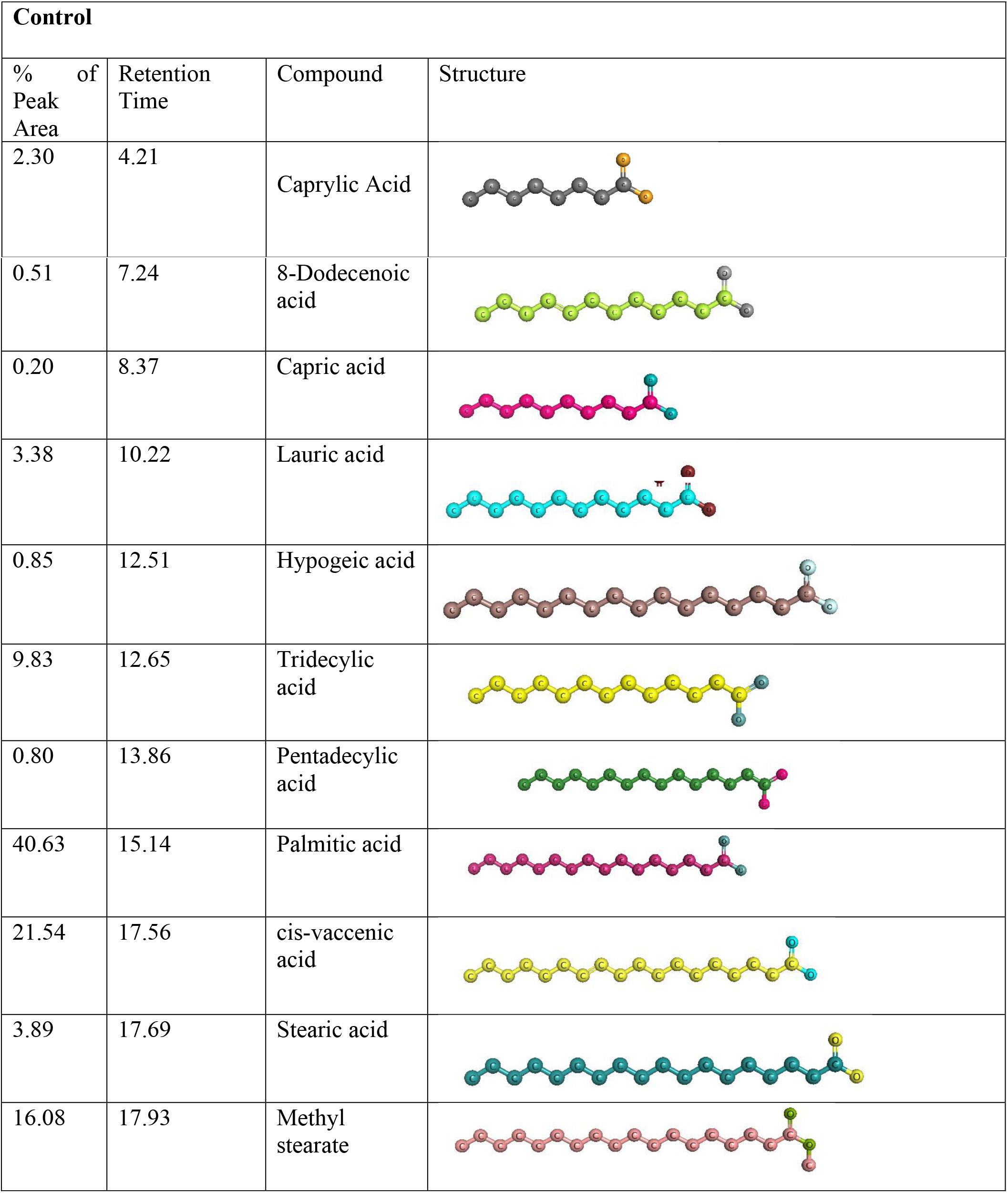
GC/MS analysis of the fatty acid profile of the control milk sample.

| **Control** |  |  |  |
| --- | --- | --- | --- |
| % of Peak Area | Retention Time | Compound | Structure |
| 2.30 | 4.21 | Caprylic Acid |  |

|  |  |  |
| --- | --- | --- |
| 0.51 | 7.24 | 8-Dodecenoic acid |
| 0.20 | 8.37 | Capric acid |
| 3.38 | 10.22 | Lauric acid |
| 0.85 | 12.51 | Hypogeic acid |
| 9.83 | 12.65 | Tridecylic acid |
| 0.80 | 13.86 | Pentadecylic acid |
| 40.63 | 15.14 | Palmitic acid |
| 21.54 | 17.56 | cis-vaccenic acid |
| 3.89 | 17.69 | Stearic acid |
| 16.08 | 17.93 | Methyl stearate |

**Table 3.**
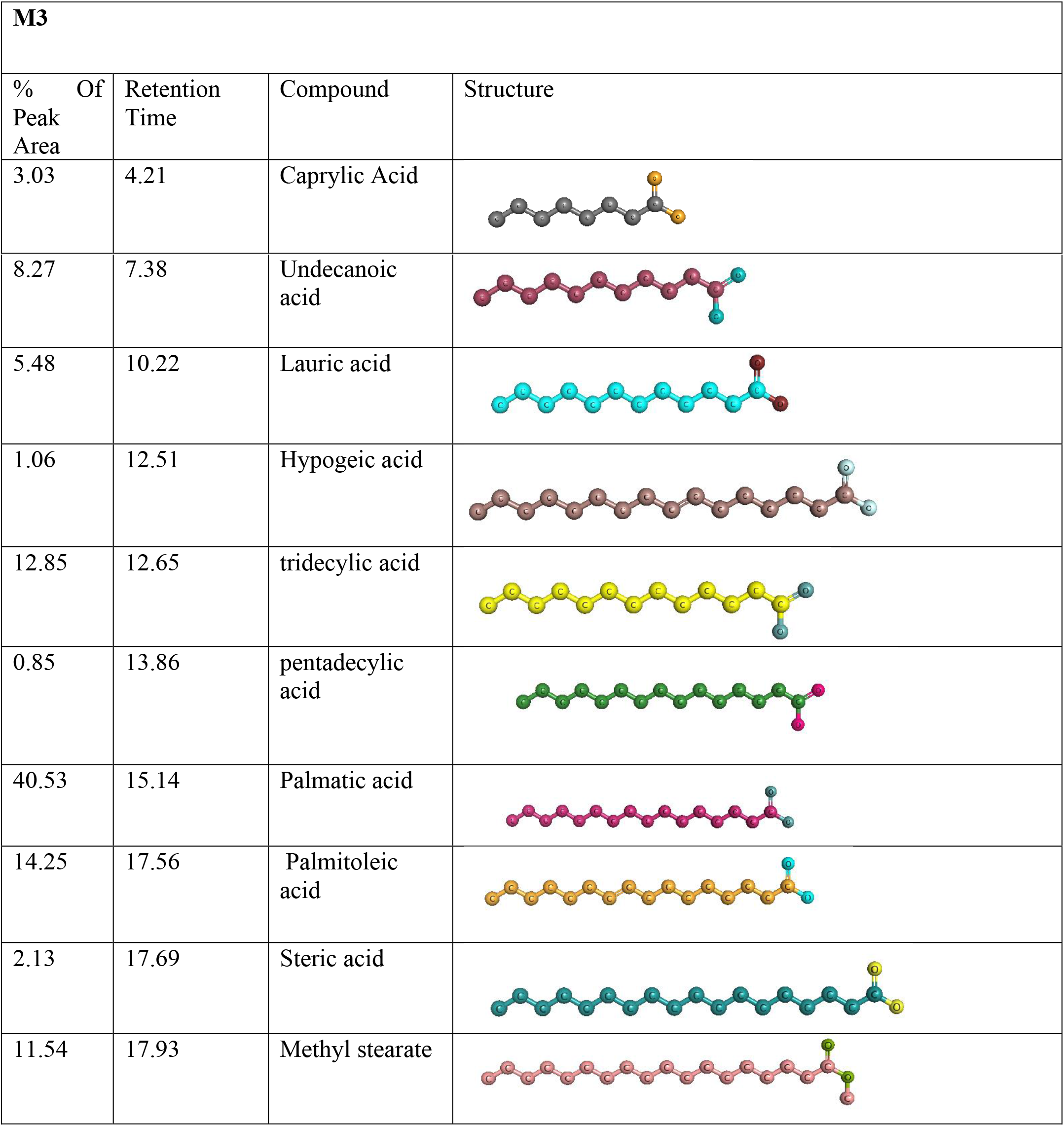
GC/MS analysis of the fatty acid profile of plasma bubbled milk sample.

| M3 |  |  |  |
| --- | --- | --- | --- |
| % Of Peak Area | Retention Time | Compound | Structure |
| 3.03 | 4.21 | Caprylic Acid |  |
| 8.27 | 7.38 | Undecanoic acid |  |
| 5.48 | 10.22 | Lauric acid |  |
| 1.06 | 12.51 | Hypogeic acid |  |
| 12.85 | 12.65 | tridecylic acid |  |
| 0.85 | 13.86 | pentadecylic acid |  |
| 40.53 | 15.14 | Palmatic acid |  |
| 14.25 | 17.56 | Palmitoleic acid |  |
| 2.13 | 17.69 | Steric acid |  |
| 11.54 | 17.93 | Methyl stearate |  |

Further, it was observed that the omega 9 fatty acid such as hypogeic acid increases having a peak area of 1.06% while in the control sample, the value of peak area was 0.85%. Again, lauric acid a saturated fatty acid was increased in post plasma treatment with a peak area of 5.48%, whereas in control the peak area was 3.38%. On the other hand, the formation of palmitoleic acid (omega 7 fatty acid) formation which proved the increase in triglycerides level (aldehydes) after exposure to plasma bubbling. So, the increase in triglyceride level (hypogeic acid, lauric acid and palmitoleic acid) could be explained due to the formation of ozone during cold plasma and further leads to decomposed and thus resulting in the formation of aldehyde (Díaz *et al*. 2003; Whitehead, 2016). Overall, a good and non-detrimental effect on the fatty acid profile was observed after exposure to plasma bubbling treatment which could be used in future in the food industry.

### Conclusion

Cow milk is a delicate liquid food; the use plasma bubbling system was able to decontaminate microbes such as coliform, yeast and mould. Sensitive parameters like pH, TSS, colour, TA and GC-MS were measured after the treatment of plasma bubbling. The validation of the optimised condition of the plasma bubbling system depends on the input processing parameters such as voltage, time interval, flow rate of air and volume of the milk sample which revealed that all these variables markedly affect the microbial load as well as physicochemical parameters of cow milk. The optimum conditions of plasma bubbling for milk were 200 V, 10 L/h, 15 min time interval with 100 mL of cow milk sample in which no viable cell of microorganism was detected and no negative effect was found in physiochemical properties of cow milk. To better understand the plasma bubbling of milk, in future the sequence analysis of protein and *in vivo* experiments should be done to observe the biological effect of cold plasma treated milk which is necessary to market the product.

## Notes

### Competing Interest Statement

The authors have declared no competing interest.

### Summary of Updates

this manuscript contains proper figure number

